# The Magnitude of Telomere Shortening per Cell Division In Vivo: Implications for Lifelong Hematopoiesis in Humans

**DOI:** 10.1101/2025.04.16.649166

**Authors:** Jennifer R Harris, Troels Steenstrup, Abraham Aviv

## Abstract

The magnitude of telomere shortening per cell division in human somatic cells in vivo (MTS_IV_) is a fundamental but unquantified parameter. MTS_IV_ is essential for understanding how telomere-length (TL)-dependent hematopoietic cell division influences age-related health and longevity. By leveraging sex differences in leukocyte TL and the differential dosage of *DKC1*, a telomerase-regulating gene, during early embryonic cell divisions, we estimate the MTS_IV_ to be 28 base pairs per cell division (95% CI: 23 – 32). Using this estimate and leukocyte TL data from newborns and centenarians, we infer that hematopoietic stem cells (HSCs) undergo approximately 156 divisions (95% CI: 130-183) over a 100-year lifespan. Using longitudinal data on leukocyte telomere shortening in adults, we further estimate that HSCs divide about 0.97 times yearly (95% CI: 0.80 - 1.13) after age 20. These findings provide a quantitative framework for understanding TL-dependent hematopoiesis, the most proliferative process in the human soma. They also highlight that if telomere shortening affects age-related health and longevity, it acts primarily through its impact on hematopoiesis. Our results refine hematopoietic stem cell replicative history estimates and might guide treatments involving hematopoietic cell expansion, such as hematopoietic cell transplantation and immunotherapies.

**Significance Statement:** When human somatic cells divide, their telomeres shorten. This process drives age-related shortening of leukocyte telomeres and reflects hematopoietic stem cell (HSC) division at the top of the hematopoietic hierarchy. The magnitude of telomere shortening per somatic cell division in vivo (MTS_IV_) is a critical yet previously unquantified parameter essential for providing insights into the pace of HSC division in humans. This study estimates that the MTS_IV_ is about 28 base pairs. Using the MTS_IV_ and leukocyte telomere length data from newborns and centenarians, we calculated that HSCs divide approximately 156 times over a 100-year lifespan. This knowledge improves our understanding of HSC replication dynamics and their implications for human health and longevity.

## Introduction

The magnitude of telomere shortening with each somatic cell division in vivo (MTS_IV_) provides a crucial quantitative link between telomere length (TL) and the body’s replicative demands. Hematopoiesis is the most proliferative TL-dependent process in the human soma (*1*). Therefore, knowledge of MTS_IV_ is particularly relevant for understanding the role of hematopoietic cell (HC) replication in aging-related diseases and longevity (*2*). Knowing the MTS_IV_ may also have important implications for the outcomes of HC transplants (1, 2) and anti-cancer immunotherapies, including tumor-infiltrating lymphocytes (TILs) (3) and chimeric antigen receptor (CAR) T-cells (4). The success of these treatments depends on the TL-dependent replicative capacity of HCs infused into recipients.

However, a reliable estimate of MTS_IV_, crucial for measuring TL-dependent replicative capacity in vivo, remains unknown. To address this gap, we leveraged sex differences in leukocyte TL (LTL) and data on delayed X-chromosome inactivation in female embryos to estimate the MTS_IV_. We use LTL because it reflects age-related shortening of telomeres in hematopoietic stem cells (HSCs) (5).

### Approach

The X-chromosome harbors *DKC1*, which encodes dyskerin, a protein essential for the activity of telomerase (6), the reverse transcriptase that elongates telomeres (7). Detrimental *DKC1* mutations cause dyskeratosis congenita, an X-linked telomere biology disorder (TBD) that presents with critically short leukocyte telomeres and bone marrow failure (8). Differential *DKC1* dosage during early embryogenesis (9) likely explains why newborn females have longer leukocyte telomeres than newborn males (10) ─ a difference that persists throughout life (10, 11).

Females inherit an X chromosome from each parent, one of which is randomly inactivated during early embryogenesis. In humans, this occurs right after the formation of the early blastocyst. The timing of X chromosome inactivation (12) enables determining the effect of the differential *DKC1* dosage between female and male embryos on TL during the early embryonic cell divisions, from the zygote to the blastocyst stage.

The zygote undergoes four cell divisions (cleavages) to form the 16-cell morula, followed by two additional divisions to form the earliest blastocyst, comprising about 64 cells (**Figure 1**) (12). Accordingly, X-chromosome inactivation in female embryos likely occurs after 6-7 cell divisions, during which X-linked genes, including *DKC1*, are expressed biallelically in female embryos. This suggests that during the first 6-7 cycles of early embryonic cell division, female embryos experience approximately double the *DKC1* dosage─ and, presumably, double telomerase activity, compared to male embryos (**Table 1, Figure 1**).

**Table 1:** Parameters, Calculated Values and Methods.

| Parameter | Description | Value | CI (95%) | Method/Reference |
| --- | --- | --- | --- | --- |
| A | Sex difference in leukocyte TL in adults (bp) | 179 | (153 - 206) | Meta-analysis based on (11) |
| B | Number of divisions before blastocyte | 6.50 | (6.00 - 7.00)* | Constructed so CI is 6-7 |
| C = A/B | MTSIV per cell division in vivo (bp) | 28 | (23 - 32) | Derived from numbers of cell divisions before blastocyte and meta-analysis based on data in (11), using Delta method. |
| D | Yearly leukocyte TL shortening (bp) | 27 | (26 - 28) | Meta-analysis of data in (17) |
| E = D/C = D x B/A | HSC divisions per year after age 20 | 0.97 | (0.80 - 1.13) | Calculated based on a meta-analysis of data in (11) and (17), and B using the delta method |
| F | Leukocyte TL at age 100 (kb) | 5.2 | (5.1 – 5.3) | Meta-analysis data in (16) |
| G | Leukocyte TL at birth (kb) | 9.5 | (9.4 – 9.6) | (10) |
| H = G – F | Leukocyte TL change from birth to 100 (kb) | 4.3 | (4.2 – 4.4) | Derived from F and G values |
| I = H/C = (G - F) x B/A | Number of HSC divisions from birth to 100 | 156 | (130- 183) | Calculated from (10) and meta-analysis of data in (16) and in (11), and derived values for C, H, and CI for B using the Delta method |

\*For the number of cell divisions before X-chromosome inactivation, we use the range of 6-7 as the 95% CI constructed symmetrically around the point estimate of 6.5 kb.
TL- telomere length; MTSIV- magnitude of telomere shortening; HSC- hematopoietic stem cell

**Figure 1.**
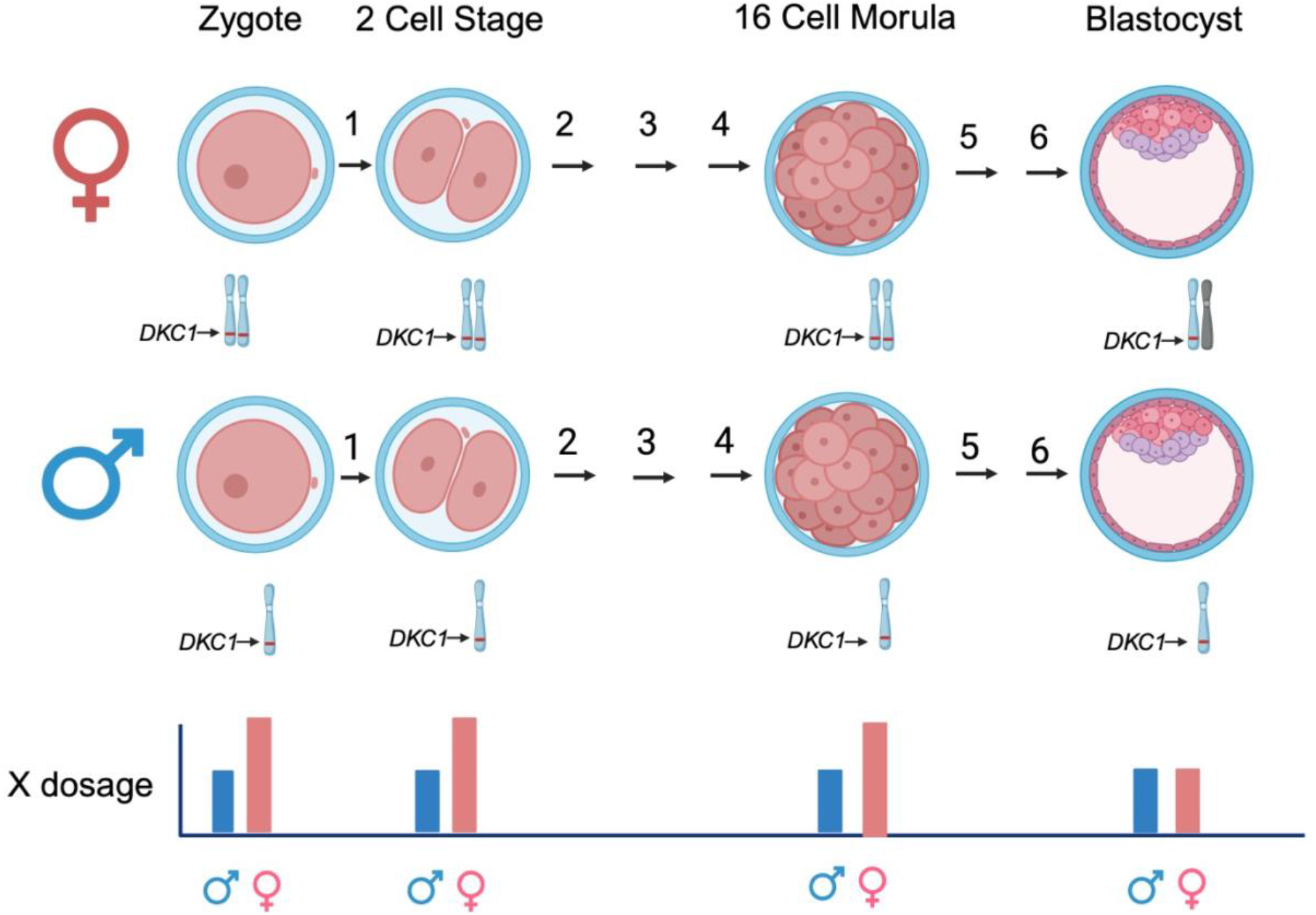
X-chromosome gene dosage (including *DKC1*) during early embryogenesis. Numbers denote cell divisions. X-chromosome inactivation presumably occurs right after early blastocyst formation, at about the 6^th^ or 7^th^ cell division. A dark X-chromosome denotes an inactivated X-chromosome. Created in https://BioRender.com.

Based on data from 12,320 adults, leukocyte TL in females averages 179 base pairs (bp) longer than that in males (**Table 1**) (11). Assuming a double-dose effect of *DKC1* during the first 6-7 cycles of embryonic cell division, female embryos lengthen their telomeres by about 28 bp per division than male embryos. (**Table 1, Figure 2**). A logical corollary is that, following X-chromosome inactivation, the single dose of *DKC1* would prevent telomere shortening by about 28 bp per division in both female and male embryos. Moreover, once telomerase is repressed postnatally (13), the MTS_IV_ would be about 28 bp in both females and males (**Figure 2**).

**Figure 2.**
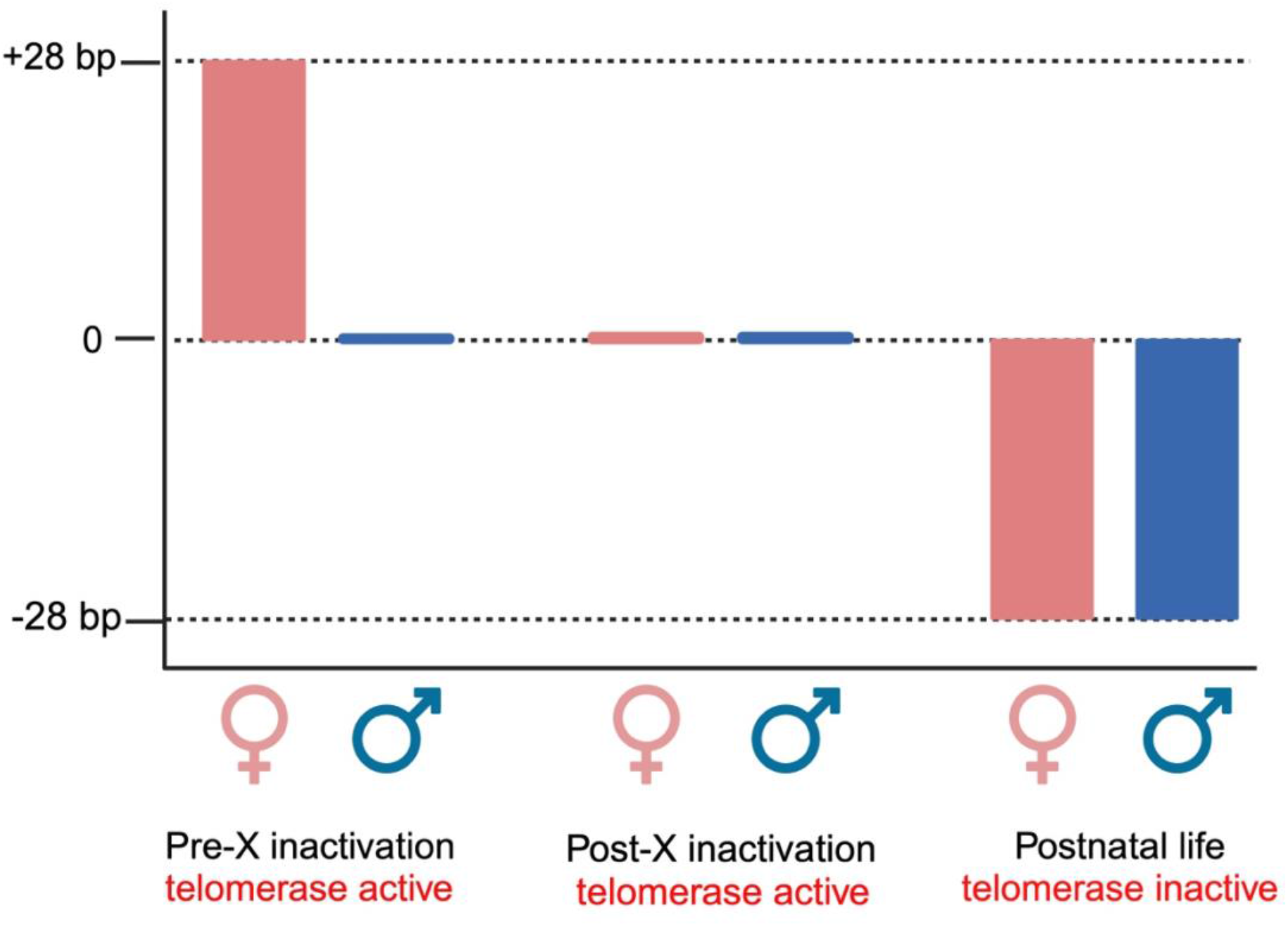
Change in telomere length per cell division in females and males. Pre-X-chromosome inactivation, female embryos experience about double-dose of *DKC1*. This results in about 28 bp gain per division in the females than males. Post-X-chromosome inactivation, all embryos are exposed to a single *DKC1* dose. Once telomerase is repressed, telomeres shorten by 28 bp with cell division in all somatic cells of both sexes. Created in https://BioRender.com

We next calculated the number of divisions HSCs undergo from birth to 100 years of age (**Table 1**). Hematopoiesis involves successive waves of cell replication, starting from the top of the hematopoietic hierarchy and progressing downward. Approximately 100,000 HSCs and multipotent progenitor cells divide at the apex (14), generating about 300-400 billion circulating blood cells released daily from the bone marrow. The age-related shortening of leukocyte telomeres thus reflects the telomere shortening in HSCs at the top of the hematopoietic hierarchy (**Figure 3**) (5, 15).

**Figure 3.**
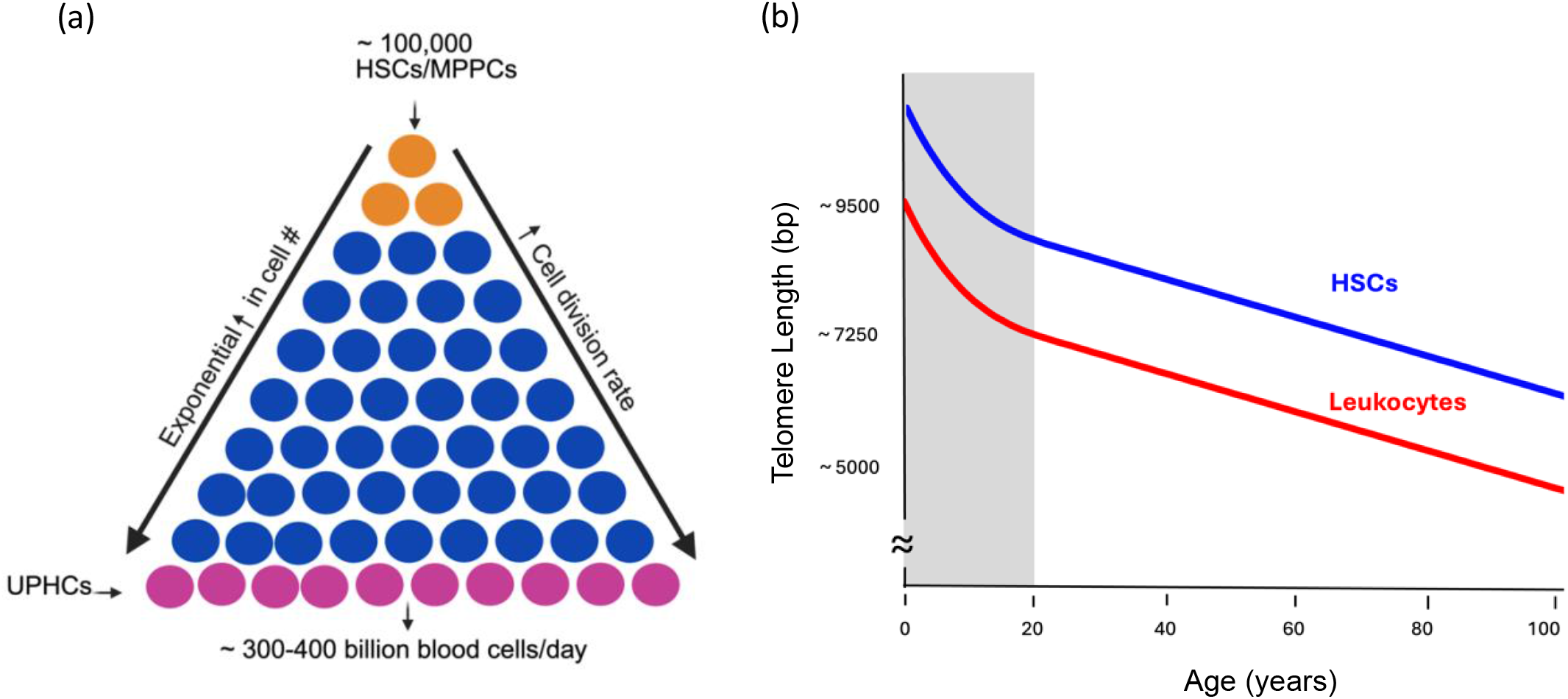
The hematopoietic hierarchy and telomere shortening in hematopoietic cells with age. (a) Around 100,000 hematopoietic stem cells (HSCs) and multipotent progenitor cells (MPPCs) at the top of the hierarchy generate oligopotent and subsequently unipotent progenitor cells (UPPCs). As cells progressively differentiate down the hierarchy, their replication rate accelerates, increasing their numbers exponentially. Approximately 300-400 billion blood cells are released daily into the circulation. (b) Leukocyte telomere shortening with age mirrors HSC shortening at the apex of the hematopoietic hierarchy. Figure 3(a) created in https://BioRender.com

Relatedly, circulating leukocytes consist of myeloid and lymphoid cells. Most myeloid cells are fully differentiated and do not replicate after leaving the bone marrow. Their age-related telomere shortening thus reflects that of HSCs in the bone marrow. In contrast, lymphocytes continue replicating in extramedullary sites, and consequently, their telomeres are shorter than those of myeloid cells. However, LTL is highly correlated with myeloid cell TL (r = 0.990) throughout the human lifetime (5), indicating that age-dependent leukocyte telomere shortening reliably reflects HSC telomere shortening.

At birth, the average LTL is 9.5 kilobases (kb) (based on data from 490 newborns (10)) and by the age of 100, it shortens to 5.2 kb (based on data from 132 centenarians (16)). This represents an average total telomere shortening of 4.3 kb over a century (**Table 1**). An MTS_IV_ of 28 bp corresponds to about 156 HSC divisions over 100 years.

We then estimated the division rate of HSCs in adults (20 or older) based on longitudinal LTL measurements from four studies involving 1,156 adults (17). After the second decade of life, leukocyte telomeres shorten by an average of 27 bp per year (**Table 1**). Applying the estimated MTS_IV_ of 28 bp, we calculated that HSCs divide 0.97 times yearly in adults. From this division rate, we estimate that of the 156 total HSC divisions occurring over a 100-year lifetime, approximately half (78 divisions) occur between ages 20 and 100, while the remaining half occur during the first two decades of life (**Figure 4**).

**Figure 4.**
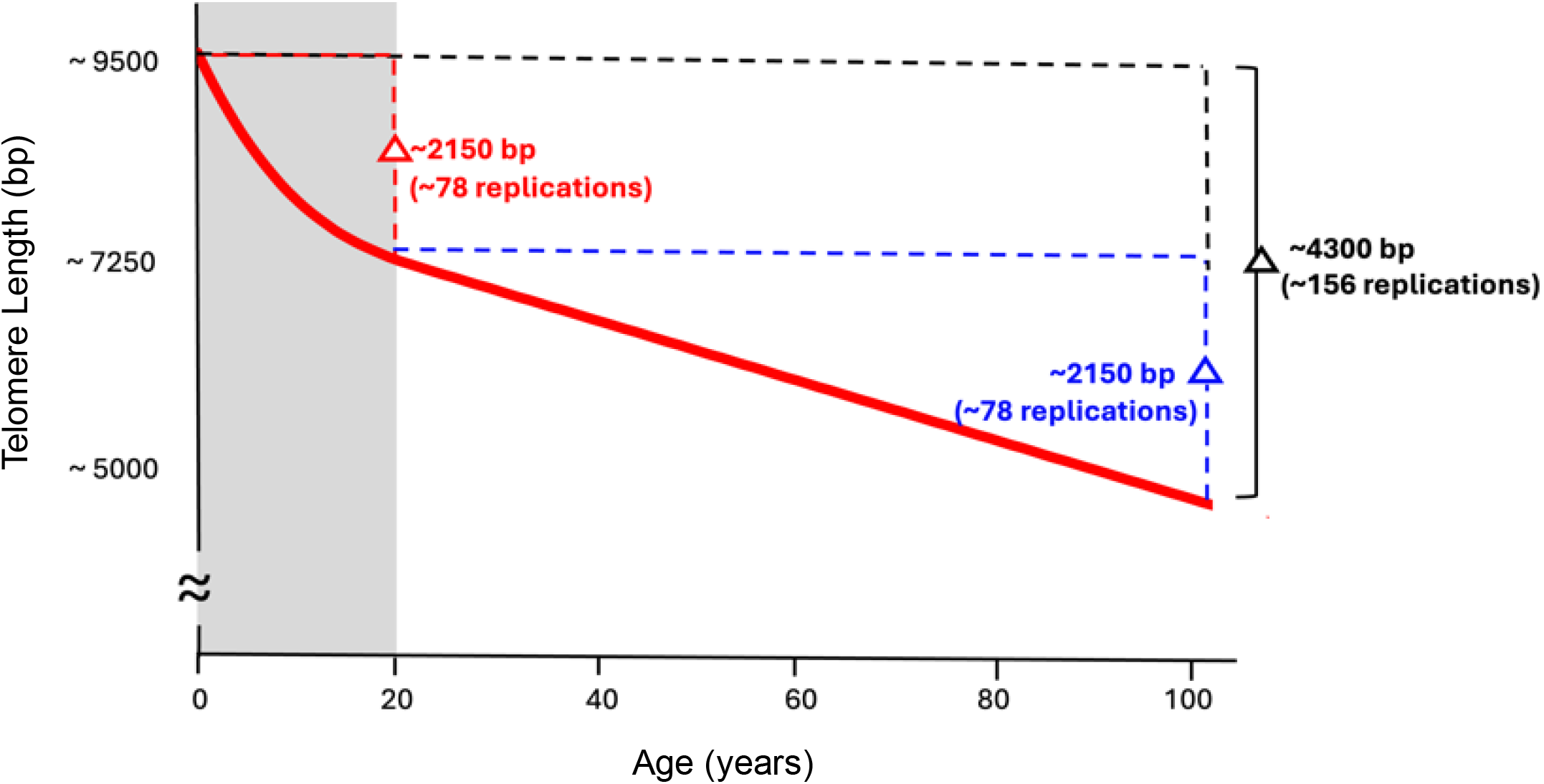
Estimating hematopoietic stem cell (HSC) replications from birth to 100 years based on leukocyte TL measurements. Assuming an MTS_IV_ of approximately 28 bp per cell division and a total leukocyte telomere shortening of ∼ 4300 bp (from ∼ 9.5 kb at birth to ∼ 5.2 kb at age 100), we estimate that HSCs replicate approximately 156 times over a human lifetime. With this MTS_IV_ estimate and analysis of published data, we calculate the rate of HSC division is about 0.97 per year after age 20, partitioning lifetime HSC replications into ∼ 78 before age 20 (gray shading) and ∼ 78 between ages 20 and 100.

## Discussion

The estimated MTS_IV_ of 28 bp advances our understanding of age-dependent leukocyte telomere shortening with HSC divisions across the human lifetime and its potential role in aging-related human diseases and longevity. This finding may also have important implications for HC transplants and therapies with immune cells if the TL-dependent division capacity of infused cells affects treatment efficacy.

The hierarchical nature of hematopoiesis has been employed in previous studies to model HSC divisions in humans. These studies analyzed age-related changes in leukocytes, focusing on either the accumulation of mutations (18) or the shift in the ratio of maternal to paternal X-chromosomes (19). They estimated that HSCs divide in vivo about once in 2–20 months (18) or 25-50 weeks (19). A more recent model based on telomere shortening with age estimated that HSCs divide about once every 1-4 years in adults and about 30-120 times over an 85-year lifetime (20). However, that estimate relies on cultured cell data showing telomere shortening of 65 (30-100) bp per cell division, which may not accurately reflect the MTS_IV_.

Our estimate that adult HSCs divide about 0.97 times a year aligns with those derived from approaches independent of TL (18, 19). This convergence of findings strengthens the validity of our estimated MTS_IV_ of approximately 28 bp per cell division. Moreover, our results, based on large sample sizes of newborns, adults, and centenarians, yielded narrower confidence intervals compared to other approaches (18-20).

The rate of HSC division—about 78 divisions in the first two decades and 78 divisions in the following eight decades—provides new insights into age-related shortening of leukocyte telomeres (21). During growth, the faster rate of telomere shortening mirrors the accelerated pace of HSC division required to support the expansion of the hematopoietic system in tandem with overall somatic growth (22). In adulthood, HSC division slows and stabilizes as the hematopoietic system transitions from expansion to maintenance.

Hematopoiesis is the most proliferative process in humans (15) and explains why leukocytes have the shortest telomeres among all adult somatic cells (23). Over a human lifetime, HC telomeres are thus likely the first among somatic cells to shorten to a critical length—a “telomeric brink”—that halts further cell replication in vivo. In this regard, approximately half of centenarians exhibit leukocyte TL comparable to that seen in patients with TBDs (11). This suggests that while MTS_IV_ of HCs and their replicative capacity play a critical role in the pathogenesis of TBDs, they may also influence health and longevity in the general population (24).

Estimates of the MTS_IV_ of HCs per cell division could also provide important information about the potential and limitations of HC transplants (1, 2) and immune cell-based therapies, such as TILs (3) and CAR-T cells (4), for treating human diseases. A study of HC transplant recipients found that better survival outcomes were associated with donor leukocyte telomeres longer than 6.7 kb. On average, recipient leukocyte telomeres shortened by 440 bp within three months post-transplant. However, survival declined sharply in recipients whose leukocyte telomeres shortened by less than 230 bp post-transplant (2). Based on MTS_IV_ of 28 bp, donated HCs underwent an average of 17 divisions in recipients, and survival decreased considerably when donated HCs underwent fewer than 8 divisions.

Similarly, an analysis of data from melanoma patients treated with TILs revealed a relationship between the TL of infused cells and treatment response, with median TL of 6.8 kb in complete responders (CR), 6.3 kb in partial responders (PR), and 4.9 kb in non-responders (NR) (3). Assuming an MTS_IV_ of 28 bp per division, the 500 bp TL difference between CR and PR amounts to about 18 additional divisions of TILs in the CR group. Similarly, the 1.9 kb TL difference between CR and NR corresponds to approximately 68 extra divisions of infused TILs in the CR group. In contrast, a small study of patients receiving CAR-T therapy for hematological malignancies (4) found no significant association between the treatment outcomes and TL in infused CAR-T cells, though that study was likely underpowered (25). To our knowledge, no other studies have examined the role of telomeres in the efficacy of these therapies, highlighting a critical research gap.

A key question is whether the estimate of MTS_IV_ varies by cell type in vivo. Culture conditions can affect telomere shortening per division in human somatic cells (26). However, consistent with the “end replication problem” (27, 28), there is no theoretical basis to support variation in telomere shortening per cell division across different somatic cell types in vivo. The slow rates of age-related telomere shortening observed in certain human tissues, such as skeletal muscle and fat, are primarily attributed to their low proliferation rates, particularly in early life (29, 30). Absent evidence showing otherwise, we assume that MTS_IV_ is about 28 bp in different somatic cells. This assumption is further supported by the consistency between our estimates of HSC replication rates with those obtained from studies that did not rely on TL (18, 19).

Finally, our estimate of MTS_IV_ is derived from multiple studies whose TL data were generated by a single laboratory. These data are based on Southern blotting (SB) of the terminal restriction fragments (TRFs) (21, 31), which, until recently, was considered the “gold standard” for TL measurements. Newly developed long-read telomere sequencing techniques (32-34) now enable the measurement of canonical TL—comprising only the TTAGGG repeats—without confounding from the sub-telomeric region, a known limitation of SB of the TRFs (21, 31). Studies leveraging this advancement in TL measurement may help further refine the MTS_IV_ estimate.

In conclusion, knowledge of the MTS_IV_ offers valuable insights into the role of HC TL in aging-related diseases and human longevity. It also can help develop metrics for determining the efficacy of HC transplants and immune-based therapies for human cancers and other diseases.

## Methods

Leukocyte TL was measured using SB of the TRFs (21, 31).

We leveraged cross-sectional leukocyte TL data from newborns (10), adults older than 20 years (11), and centenarians (16) and data on longitudinal leukocyte telomere shortening with age in adults (17). Leukocyte TL was found to be longer by 179 (95% CI: 153-206) bp in 8588 women than in 4712 men (11) and by 144 (95% CI:20 - 268) bp longer LTL in 216 newborn girls than in 274 boys (10). The sex difference in leukocyte TL between adults and newborns was insignificant (p=0.55). We thus used the adult estimate of the sex effect in our modeling (**Table 1**) since it is based on a large sample size and statistical testing confirming the consistency of the sex effect between newborns and adults.

We employed two methods for CI estimation. First, using meta-analysis, we combined data from independent studies to generate narrower CIs (35). This included: a) 95% CIs for the sex difference in leukocyte TL (11), b) leukocyte telomere shortening per year (17), and c) leukocyte TL at birth and 100 years of age (16). Second, the Delta Method (36), assuming zero covariance, was used to estimate the 95% CIs for a) the MTS_IV_ per cell division in vivo, based on sex differences in leukocyte TL (10, 11) divided by the number of embryonic cell divisions, b) the number of HSC replications per year in adults, calculated by dividing the annual leukocyte telomere shortening (17) by the MTS_IV_ per cell division in vivo, and c) the total number of replications from birth to age 100, determined by dividing the leukocyte telomere shortening from age 0 (9) to 100 years (16) by the MTS_IV_. In the text, we present data as mean values, whereas in **Table 1**, we present data as mean values with 95% CIs.

## Acknowledgments

JRH contributed extensively to the research and writing of this paper as part of her sabbatical with AA. TS is a former post-doctoral fellow of AA. The work of JRH and AA is supported, in part, by the Research Council of Norway through its Centres of Excellence funding scheme, project number 262700. AA is a member of the NIH-funded Telomere Research Network (TRN), and his recent research has also been supported by National Institutes of Health grants U01AG066529 and R01ES035760.

## Data sharing plans (for all data, documentation, and code used in analysis)

This research is based upon secondary analysis of published data as referenced in **Table 1**. Original sources and equations used to derive values, parameter estimates, and their respective confidence intervals are also presented in **Table 1**.

